# Cycloheximide-Producing *Streptomyces* Associated with *Xyleborinus saxesenii* and *Xyleborus affinis* Fungus-Farming Ambrosia Beetles

**DOI:** 10.1101/511493

**Authors:** Kirk J. Grubbs, Frank Surup, Peter H. W. Biedermann, Bradon R. McDonald, Jonathan Klassen, Caitlin M. Carlson, Jon Clardy, Cameron R. Currie

## Abstract

Symbiotic microbes help a myriad of insects acquire nutrients. Recent work suggests that insects also frequently associate with actinobacterial symbionts that produce molecules to help defend against parasites and predators. Here we explore a potential association between Actinobacteria and two species of fungus-farming ambrosia beetles, *Xyleborinus saxesenii* and *Xyleborus affinis*. We isolated and identified actinobacterial and fungal symbionts from laboratory reared nests, and characterized small molecules produced by the putative actinobacterial symbionts. One 16S rRNA phylotype of *Streptomyces* (XylebKG-1) was abundantly and consistently isolated from the nests and adults of *X. saxesenii* and *X. affinis* nests. In addition to *Raffaelea sulphurea*, the symbiont that *X. saxesenii* cultivates, we also repeatedly isolated a strain of *Nectria* sp. that is an antagonist of this mutualism. Inhibition bioassays between *S. griseus* XylebKG-1 and the fungal symbionts from *X. saxesenii* revealed strong inhibitory activity of the actinobacterium towards the fungal antagonist *Nectria* sp. but not the fungal mutualist *R. sulphurea*. Bioassay guided HPLC fractionation of *S. griseus* XylebKG-1 culture extracts, followed by NMR and mass spectrometry identified cycloheximide as the compound responsible for the observed growth inhibition. A biosynthetic gene cluster putatively encoding cycloheximide was also identified in *S. griseus* XylebKG-1. The consistent isolation of a single 16S phylotype of *Streptomyces* from two species of ambrosia beetles, and our finding that a representative isolate of this phylotype produces cycloheximide, which inhibits a parasite of the system but not the cultivated fungus, suggests that these actinobacteria may play defensive roles within these systems.

## Introduction

Ambrosia beetles are a diverse group of insects (~3,400 species) that cultivate fungi for food [1, 2]. Adult beetles generally bore into dead or dying trees, establishing a nest in the xylem. They actively inoculate the tunnel walls of the nest with spores of their mutualistic fungus, which grows and forms a layer of nutrient rich aleurioconidia (“ambrosial growth” [3]) on the woody tissue of the host plant and serves as the sole source of nutrition for adults and developing beetle larvae. Ambrosia beetles vector their fungal mutualist in specialized structures called mycangia or mycetangia [4, 5]. Nutritional symbioses with fungi evolved at least eleven times independently in bark- and ambrosia beetles (*Scolytinae* and *Platypodinae*: Coleoptera) [1, 6]. Specific ambrosia beetle species associate with specific ambrosia fungi [3, 7–9], although some beetles appear to rely on a community of cultivars. Fungal cultivars from the scolytine weevil genera *Xyleborus* and *Xyleborinus* are mostly in the Ascomycota genera *Ambrosiella* (Ceratocystidaceae: Microascales) and *Raffaelea* (Ophiostomataceae: Ophiostomatales), which convergently evolved as beetle cultivars 30-60 million years ago [10]. Whereas many phloem-boring bark beetles gain extra nutrition by associations with their cultivar fungi (e.g. *Dendroctonus* sp.), those xylem-boring ambrosia beetles that we studied are true fungus-farmers and obligately rely on their cultivars for food [9, 11]. *Nectria, Penicillium* and *Aspergillus* species are common associates of these beetles, but are typically found at low abundances within nests. They are regarded competitors, parasites or pathogens of the ambrosia beetle mutualism [3, 9].

In addition to ambrosia beetles, active farming of fungi also occurs in attine ants and macrotermitine termites [12–14], and nutritional symbioses with fungi are widespread in insects [15–21]. Reliance on fungi by these insects exposes them to potential parasite pressure in the form of pathogens or competitors of their symbionts. For example, the fungal mutualist of attine ants is impacted by a specialized and potentially virulent fungal parasite [22, 23]. To help defend the cultivar from this parasite the ants use actinobacterial symbionts that produce antibiotics [23–26]. A similar type of defensive symbiosis has been shown in the fungus-associated bark beetle *Dendroctonus frontalis* [27], and has been further suggested in the Mediterranean Pine Engraver bark beetle, *Orthotomicus erosus* [28], as well as fungus-growing termites [29]. Beyond defending fungal mutualists in agricultural associations, Actinobacteria are well adapted for insect dispersal (e.g. by desiccation-resistant, hydrophobic spores that stick to the surface of insects [30]) and fulfill different defensive capacities in other insect systems. Within antennal glands, Beewolves (*Philanthus* spp.) cultivate Actinobacteria that they transfer into brood cells and onto developing cocoons in order to prevent infection by a wide range of pathogens [31]. Actinobacteria and the antibiotic secondary metabolites they produce have been identified in several species of mud daubers [32, 33]. Furthermore, Actinobacteria have been isolated from several additional ant species [34–36] and the gypsy moth [34].

The majority of insect defensive symbioses characterized have involved Actinobacteria, which is not surprising as Actinobacteria, especially *Streptomyces*, are well known producers of bioactive secondary metabolites [37]. Over 10,000 biologically active compounds have been identified from Actinobacteria, accounting for ~45% of known microbial metabolites [38]. The phylum Actinobacteria is composed of Gram-positive bacteria and is one of the largest in the domain Bacteria. They are common soil microbes, and recent studies have also identified them as dominant community members in both freshwater [39] and marine [40] habitats. As such, Actinobacteria are common microbiome constituents in many environments.

The fruit-tree pinhole borer *Xyleborinus saxesenii* Ratzeburg and the sugarcane shot-hole borer *Xyleborus affinis* Eichhoff colonize a wide variety of dying or recently dead tree species and are two of the most widespread ambrosia beetles worldwide [41, 42]. Both species are facultatively eusocial depending on the viability of the wood resource and may settle the same nest for multiple generations: Adult offspring of a single, sib-mated foundress typically delay dispersal from their mothers’ tunnel system and help her with nest-hygiene, brood-care and fungus-farming [43–45]. Unique for Holometabola, ambrosia-beetle larvae also help in these cooperative tasks [46]. The beetles’ activity and presence is necessary to maintain the fruiting and monocultures of their fungal cultivars [2, 3]. Both species are obligately dependent on *Raffaelea* ambrosia fungi [9, 47]. Experiments in *X. saxesenii* showed that these cultures are protected against pathogenic fungi, such as *Paecilomyces variottii* and *Fusarium merismoides*, by larvae and adults in unknown ways [46] and it is possible that this defence involves “microbial helpers”.

Here we describe actinobacterial symbionts of *X. saxesenii* and *X. affinis* ambrosia beetles and explore their potential function in helping defend nests against an antagonistic fungus that was isolated from *X. saxesenii*. Using specific media, we isolated both Actinobacteria and fungi from laboratory reared nests. Actinobacterial isolates were characterized using 16S rRNA gene sequencing and tested for their ability to inhibit the growth of both mutualistic and parasitic fungal isolates from the same nests. Active compounds were isolated using bioassay-guided HPLC fractionation, chemically characterized using NMR spectroscopy and mass spectrometry, and further tested using bioassays to confirm growth inhibition activity. We sequenced the genome of one actinobacterial isolate [48] to confirm this strains' phylogenetic identification, and identified a putative biosynthetic gene cluster for one of the characterized active compounds. Based on these results, we propose a mutualism between two species of ambrosia beetle and Actinobacteria, in which the bacterial symbiont produces cycloheximide to inhibit the growth of fungal competitors of the mutualistic cultivar fungus.

## Materials and Methods

### Beetle collection and rearing

*Xyleborus affinis* and *Xyleborinus saxesenii* females (N > 20 each) were collected at the Southern Research Station in Pineville, LA (31°20’ N, 92°24’ W; 123 ft asl) with four ethanol (95%) baited Lindgren funnel traps in October 2007. Live beetles were placed in sterile plastic tubes with wet filter paper, stored at 4 °C for up to three days, surface sterilized by immersing in 70% ethanol and deionized water for a few seconds, and then reared on artificial medium in glass tubes following Biedermann *et al* [44, 49]. Briefly, beetles were reared in sterile glass tubes (Bellco culture tubes 18 × 150 mm) filled with the standard medium for rearing xyleborine ambrosia beetles. A single female per glass tube was put onto the medium and usually started boring tunnels as if in wood (N = 20 tubes/species). About one third of these beetle colonies successfully established brood and these were maintained in the lab at room temperature with indirect sunlight.

### Isolation of Actinobacteria

We conducted targeted isolation of Actinobacteria from each of three *X. saxesenii* and *X. affinis* colonies in triplicate, aseptically sampling each tube three times in a biosafety cabinet. Briefly, the nest inside the solid rearing substrate was shaken out of the tube and tunnel-wall material containing the layer of the mutualistic fungus (henceforth termed nest material), as well as individuals were collected with sterile metal probes / tweezers from the exposed tunnels. *X. saxesenii* nest material (0.05 g per sample), adults (2 pooled individuals per sample), and larvae (5 pooled individuals per sample) were sampled; only nest material (0.05 g per sample) and adults (2 pooled individuals per sample) were sampled from *X. affinis*. All samples were chosen at random and homogenized in 500 μL of autoclaved, 0.22 μm filtered, deionized water; 100 μL of each was evenly spread on dried chitin agar plates (15 g agar, 3 g chitin, 0.575 g K_2_HPO_4_, 0.375 g MgSO_4_ x 7H_2_O, 0.275 g KH_2_PO_4_, 0.0075 g FeSO_4_ × 7H_2_O, 0.00075 g MnCl_2_ × 4H_2_O, and 0.00075 g ZnSO_4_ × 7H_2_O dissolved in 750 mL deionized water) in duplicate and allowed to dry before wrapping with parafilm. Plates were incubated at 30 °C for three weeks, after which colony forming units (CFUs) were counted and eight of each morphotype per plate were transferred to yeast malt extract agar (YMEA: 4 g yeast extract, 10 g malt extract, 4 g dextrose, and 15 g agar dissolved in 1 L). Colonies on YMEA plates were allowed to grow at 30 °C for two weeks, visually inspected for morphological properties characteristic of Actinobacteria, and sub-cultured as necessary to obtain pure cultures. Three 0.05 g samples of artificial medium from tubes that were not inoculated with beetles were also plated in duplicate on YMEA without antibiotics to screen for contamination and possible presence of Actinobacteria in the beetle medium. All media used for actinobacterial isolation had filter-sterilized cycloheximide (0.05 g/L) and nystatin (10,000 units/mL) added after autoclaving and cooling to suppress fungal growth.

### Fungal isolations

Fungal symbionts were isolated from three *X. saxesenii* nests, sampled three times each. *X. affinis* were not sampled for fungi. Nest material was scraped using a sterile metal probe and inoculated on potato dextrose agar plates (PDA; Difco, Sparks, MD) with penicillin (0.05 g/L) and streptomycin (0.05 g/L) added after autoclaving and cooling to suppress bacterial growth, and incubated at 30 °C for one week. During incubation, fast growing fungi were sub-cultured onto fresh PDA plates and the agar on which they grew was fully removed to prevent overgrowth of the entire original isolation plate. Two different fungi were obtained in pure culture by successive rounds of scraping a small amount of material from the edge of each colony and then plating on individual PDA plates.

### DNA sequencing

The 16S rRNA gene was sequenced from eight Actinobacteria isolates obtained from both *X. saxesenii* and *X. affinis* for a total of 16. In an effort to maximize the possibility of capturing any phylogenetic diversity, and thereby discover if multiple species were present, the strains that were sequenced were chosen based on morphological differences rather than origin. Only two morphologies were observed with the only differences being that the spores of one morphology were slightly darker than the other. The 16S rRNA gene PCR primers used were the Actinobacteria-specific F243 (5'-GGATGAGCCCGCGGCCTA-3') and R1378 (5'-CGGTGTGTACAAGGCCCGGGAACG-3') [53], and in separate reactions the general bacterial primers pA (5'-AGAGTTTGATCCTGGCTCAG-3') and pH (5'-AAGGAGGTGATCCAGCCCGCA-3') to increase coverage length [54]. The cycle parameters used for each primer set was similar to those above except the annealing temperatures were 58 °C and 54 °C, respectively, and the elongation time was 95 s for primers pA and pH. Each PCR reaction was composed of 12.5 μl GoTaq (Promega), 1 μl of template DNA, and 40 μM of each primer in a final volume of 25 μl. The EF-α and 18S rRNA genes were sequenced for two each of the isolated *Raffaelea sulphurea* and the putative antagonistic fungus *Nectria* spp. DNA was extracted as previously described [50]. PCR primers NS1 (5'-GTAGTCATATGCTTGTCTC-3') and NS4 (5'-CTTCCGTCAATTCCTTTAAG-3') were used to amplify the 18S rRNA gene [51], using thermocycling parameters: 95 °C for 2 min, 35 cycles of 95 °C for 45 s, 42 °C for 45 s, 72 °C for 90 s, 72 °C for 5 min and hold at 4°C. EF-α gene PCR primers 983F (5′-GCYCCYGGHCAYCGTGAYTTYAT) and 2218R (5′-ATGACACCRACRGCRACRGTYTG) [52] were used with similar cycling parameters, except annealing temperature and elongation time were 55 °C and 130 s.

PCR amplicons were purified by adding 0.8 μl ExoSap-IT (USB) to 2 μl of PCR product diluted in 5.25 μl of autoclaved deionized water and incubating this mixture at 37 °C for 15 min and then at 80 °C for 15 min. Sanger sequencing reactions contained: 1 μl BigDye Terminator v. 3.1 (Applied Biosystems), 1.5 μl Big Dye Buffer (Applied Biosystems) and 1 μl of 10 μM primer, and the entire cleaned amplicon solution. Sequencing PCR conditions were 95 °C for 3 min, 35 cycles of 95 °C for 20 s, 45 °C for 30, 60 °C for 4min, 72 °C for 7 min and hold at 4°C. Excess dye terminators were removed using CleanSeq beads (Agencourt Biosciences) and samples were resuspended in 40 μl of sterile ddH_2_O and sequenced at the University of Wisconsin-Madison Biotechnology Center using an ABI 377 instrument (Applied Biosystems).

### Actinobacteria antifungal bioassays

Growth inhibition assays were conducted between one *S. griseus* XylebKG-1 like strain (see results) isolated from each of the three *X. saxesenii* nests and both isolated fungal species by first inoculating the Actinobacterium in the center of a PDA plate and allowing it to grow for two weeks. A small amount of test fungus was then inoculated at the edge of this Petri plate and grown at 30 °C for two weeks, after which a zone of inhibition (ZOI) was determined by measuring the shortest distance between the bacterium and the fungus.

### Phylogenetic analyses

All sequences were assembled using Bionumerics v6.5 (Applied Maths), searched against the GenBank Nucleotide Sequence Database [55] using BLAST [56] to determine a preliminary identity, and then aligned in MEGA5 [57] using MUSCLE [58]. 18S rRNA and EF-α sequences were aligned and trimmed individually and subsequently concatenated to increase phylogenetic resolving power. To ensure codons were not split by gaps, alignments were inspected in MEGA5 for consistent reading frames. Substitution models were chosen using the model selection module of MEGA5. Maximum likelihood phylogenies were inferred using 500 bootstrap replicates using MEGA5.

### Genome Based Phylogeny

The genome of *S. griseus* XylebKG-1 has previously been sequenced [48] allowing us to generate a genome based phylogeny for this isolate. Proteins from all complete *Streptomyces* genomes were predicted using prodigal [59] for consistency and annotated using HMMer [60] models generated from KEGG [61] gene families, of which 1,364 KEGG gene families were conserved in all genomes. For these gene families, the proteins with the highest HMMer bitscore from each genome were aligned using MAFFT [62] and then converted to a nucleotide alignment. These alignments were concatenated and a phylogeny generated using RAxML [63] with 100 rapid bootstraps.

### Synteny Map

The genomes of *Streptomyces griseus* subsp. *griseus* NBRC13350 [NC_010572.1] and *Streptomyces griseus* XylebKG-1 were aligned using progressive Mauve [64] with default parameters.

### Analytical Chemistry Methods and Instrumentation

One- and two-dimensional NMR spectra were acquired using a Varian Inova spectrometer with a frequency of 600 MHz for ^1^H and 150 MHz for ^13^C nuclei. All compounds were dissolved in CD_3_OD. HPLC/MS analysis was performed on an Agilent 1200 Series HPLC / 6130 Series mass spectrometer. High resolution spectra were obtained on a Waters Micromass Q-TOF Ultima ESI-TOF mass spectrometer.

### Isolation and Elucidation of Bioactive Compounds

*S. griseus* XylebKG-1 was cultivated on PDA plates for 5-10 days. Seed biomass for 1 L cultures was produced by adding 1 cm^2^ of a single mature PDA culture to three 500 mL Erlenmeyer flasks containing 85 mL modified yeast peptone maltose medium (YPM: 2 g/L yeast extract, 2 g/L bactopeptone, 4 g/L D-mannitol). These were incubated at 28 °C with shaking at 250 rpm for 48 h. Twenty-five ml of each culture was added to eight 1 L of YPM in 4 L Erlenmeyer flasks and incubated for 72 h at 28 °C with shaking at 250 rpm. Supernatants and mycelia were processed separately after cultures were centrifuged at 7000 rpm for 30 min. Culture supernatants were adjusted to pH 6 and extracted twice with an equal volume of ethyl acetate. After evaporation *in vacuo*, residues were resuspended in 2 mL MeOH/H_2_O (8:2). Mycelia were lyophilized and each extracted with 50 mL acetone and 50 mL methanol. After evaporation *in vacuo*, crude extracts were resuspended in 2 mL methanol. Crude supernatant and mycelium extracts were tested for inhibition of *Nectria* sp.; only the extracts of the crude supernatant showed significant assay activity. Crude supernatant extracts were purified using a 2 g pre-packed C_18_ Sep-Pak resin and fractionated by eluting with a gradient of pure water to pure methanol. The pure water flow through and 10% methanol fractions exhibited the highest anti-*Nectria* activity. These fractions were therefore combined and fractionated by gel chromatography using Sephadex LH-20 with methanol as the mobile phase (column 60 x 2.5 cm). Active fractions were combined and subsequently purified by reversed-phase HPLC (Agilent 1100 Series HPLC system, Supelco Discovery HS C18 column, 250 × 10 mm, 2 mL/min). HPLC conditions used: 2 min 80% A, 20% B in 28 min to 100% B (A: water, B: methanol). The fraction most active against *Nectria* sp. was eluted from 17.5 and 18 min and contained 2.7 mg of cycloheximide (**1**).

### One Strain Many Compounds (OSMAC) Screening

*S. griseus* XylebKG-1 strain was cultivated on agar plates (300 mL) of YPM, PDA, oat media (20 g/L oat meal, 2.5 mL/L trace element solution, 3 g/L CaCl_2_·2 H_2_O, 1 g/L Fe(III)-citrate, 0.2 g/L MnSO_4_, 0.1 g/L ZnCl_2_, 25 mg/L CuSO_4_·5 H_2_O, 20 mg/L Na_2_B_4_O_7_·10 H_2_O, 4 mg/L CoCl_2_, 10 mg/L N_a2_M_o_O_4_·2 H_2_O), soy mannitol media (20 g/L soy meal, 20 g/L mannitol), starch-glucose-glycerol media (10 g/L glucose, 10 g/L glycerol, 10 g/L starch, 2.5 mL/L cornsteep liquor, 5 g/L casein-peptone, 2 g/L yeast extract, 1 g/L NaCl, 3 g/L CaCO_3_), ISP1 media (5 g/L pancreatic digest of casein, yeast extract 3 g/L), ISP2 media (4 g/L yeast extract, 10 g/L malt extract, 4 g/L dextrose), 1187 media 10 g/L starch, 2 g/L (NH_4_)_2_SO_4_, 1 g K_2_HPO_4_, 1 g/L MgSO_4_·7 H_2_O, 1 g/L NaCl, 2 g CaCO_3_, 5 mL/L trace element solution) for 7 days at 30 °C. All plates were extracted with ethyl acetate. Naramycin B (0.7 mg, **2**) was isolated from the crude extract of ISP1 cultivation by semipreparative HPLC (gradient 22% to 40% acetonitrile in 25 min, Supelco Discovery HS C18 column, 250 × 10 mm). Actiphenol (1.6 mg, **3**) was isolated from the crude extract of 1187 cultivation by preparative HPLC (gradient 65% to 100% methanol in 25 min, column Phenomenex Luna C18 250×21 mm. Dihydomaltophilin (2.4 mg, **4**) was isolated from extracts of PDA and 1187 cultivations by preparative HPLC (gradient 50% to 100% acetonitrile in 25 min, column Phenomenex Luna C18 250×21mm), (gradient 80% to 100% acetonitrile in 25 min, column Phenomenex Luna C18 250×21mm) and semi-preparative HPLC (gradient 35% to 50% acetonitrile in 25 min, column Supelco C18 250×8mm).

Cycloheximide (**1**): white amorphous powder; ^1^H, ^13^C NMR were identical to a commercial sample obtained from Sigma-Aldrich [65], ESI-MS *m/z* [M+Na]^+^ 304.1, [M+H]^+^ 282.1, [M−H]^−^ 280.1; HR-ESI-MS *m/z* 282.1718 [M+H]+ (calculated for C_15_H_24_NO_4_, 282.1700). Naramycin B (**2**): white amorphous powder; 1H NMR data were consistent with those previously published for this metabolite [66]; ESI-MS *m/z* [M+Na]^+^ 304.1, [M+H]^+^ 282.1, [M−H]^−^ 280.1. Actiphenol (**3**): white amorphous powder; ^1^H NMR data were consistent with those previously reported [67]; ESI-MS *m/z* [M+Na]^+^ 298.0, [M+H]^+^ 276.2, [M-H]^−^ 274.1. Dihydromaltophilin (**4**): white amorphous powder; ^1^H NMR and ^13^C NMR data were consistent with those previously reported [68]; ESI-MS *m/z* 513.3 [M+H]^+^, 511.3 [M−H]^−^; HR-ESI-MS 513.2964 [M+H]+ (calculated for C_29_H_41_N_2_O_6_ 513.2965).

### Cycloheximide antifungal assays

Minimum inhibitory concentrations were determined using *Nectria* sp. and *R. sulphurea* grown in liquid YPM for three days. Cultures were diluted 1:1000 with fresh YPM and 200 μL per well transferred into 96-well plates containing various amounts of commercial cycloheximide (100 μg, 50 μg, 20 μg, 10 μg, 5 μg, 2 μg, 1 μg) and dihydromaltophilin (5 μg, 2 μg, 0.5 μg, 0.2 μg). These 96-well plates were incubated for 72h at 30 °C, after which the optical density was measured at 600 nm using a SpectraMax M5^®^ Plate Reader. Naramycin B (**2**) and actiphenol (**3**) were inactive against both fungi up to concentrations of 10 μg/200 μL.

To determine the antifungal activity of cycloheximide (**1**) and dihydromaltophilin (**4**) in an agar plate dilution assay, *Nectria* sp. and *R. sulphurea* were grown in 20 mL liquid potato dextrose media for 7 and 21 days, respectively, at 30 °C while shaking at 250 rpm. One mL of each culture was used to inoculate PDA plates. Paper disks (6 mm diameter) were soaked with solutions of 30 μL, 2 μL, 0.2 μL of **1** and 20 μL, 2 μL of **4**, both in methanol (concentration 1 mg/mL), dried, and applied to the surface of the agar plates. Plates were grown at 30 °C for 5-7 days, when inhibition zones were recorded.

### Cycloheximide biosynthetic cluster identification

The biosynthetic cluster in the high quality draft *S. griseus* XylebKG-1 genome sequence [NZ_ADFC00000000.2] was predicted using antiSMASH v2.0 [69] and specific comparison to the previously published cycloheximide biosynthesis gene cluster from *Streptomyces* sp. YIM56141 [70]. Gene cluster functional annotations were derived from the antiSMASH output, homologous annotations in the *Streptomyces* sp. cycloheximide biosynthesis gene cluster, and retrobiosynthetic logic.

## Results and Discussion

### Isolation and identification of Actinobacteria

At least one CFU having a morphology consistent with Actinobacteria was observed from 72% of adults, 33% of larvae, and 61% of nest samples from *X. saxesenii* (Table 1). On average more Actinobacteria were cultured from nests than adults or larvae, with mean ± SEM CFUs/sample of 110 ± 56, 3.4 ± 1.4, and 1.1 ± 0.6, respectively (Fig. 1: Single factor ANOVA P = 0.0582). No growth of Actinobacteria was observed from media not inoculated with beetles (Table 1, Fig. 1). Thus, the medium serving as a possible source of bacterial isolates can be eliminated. All isolates had similar morphologies and growth patterns. Isolations from *X. affinis* nests and adults also resulted in CFUs of a single actinobacterial morphotype similar to that isolated from *X. saxesenii*.

**Table 1.** Number of plates yielding actinobacterial growth.

| *Xyleborinus saxesenii* | Adults | Larvae | Nest | Substrate |
| --- | --- | --- | --- | --- |
| Nest 1 | 6/6 | 2/6 | 6/6 |  |
| Nest 2 | 3/6 | 3/6 | 2/6 |  |
| Nest 3 | 4/6 | 1/6 | 3/6 |  |
| Control |  |  |  | 0/6 |
| Total | 13/18 | 6/18 | 11/18 | 0/6 |

**Fig. 1.**
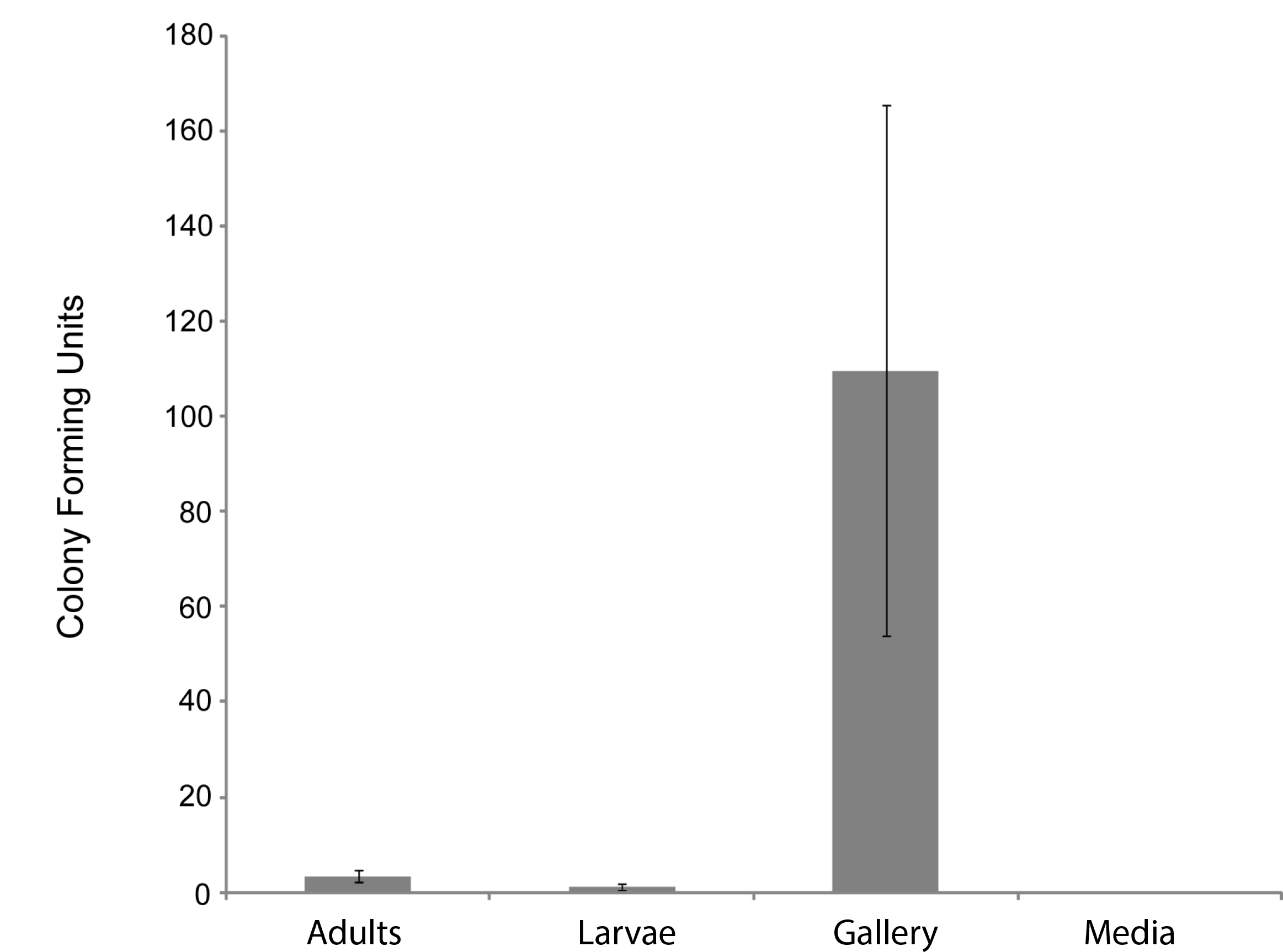
Number of Actinobacteria cultured from components of *X. saxesenii* nests. Means +/− standard errors of the mean are displayed. See methods for culture conditions.

Eight representative Actinobacteria from various samples of the *X. saxesenii* and *X. affinis* systems were identified using 16S rRNA gene sequencing. The 16S rRNA gene sequences (1,123 bp) from all 16 were 100% identical and were most similar to that of *Streptomyces griseus* subsp. *griseus* NBRC13350 [NC_010572.1] when BLAST searched against the NCBI nr database, a result confirmed by phylogenetic analysis (Fig. 2, supplemental Fig. 1). A phylogeny constructed using the genome sequence for one of these strains, *S. griseus* XylebKG-1 (XylebKG-1) isolated from *X. saxesenii* [NZ_ADFC00000000.2] confirmed the close relationship between this strain and *Streptomyces griseus* subsp. *griseus* NBRC13350 [NC_010572.1], generating a tight clade in all bootstrap replicates produced (Fig. 3). This is consistent with their high genomic similarity suggested previously using average nucleotide identity [48]. Note that although the progressive Mauve algorithm aligns both genomes as one homology block (except for the extreme 5' and 3' ends), this block contains some regions of negligible sequence homology. These regions typically represent secondary metabolite biosynthetic gene clusters of unknown function that are not conserved between these two genomes (data not shown).

**Fig. 2.**
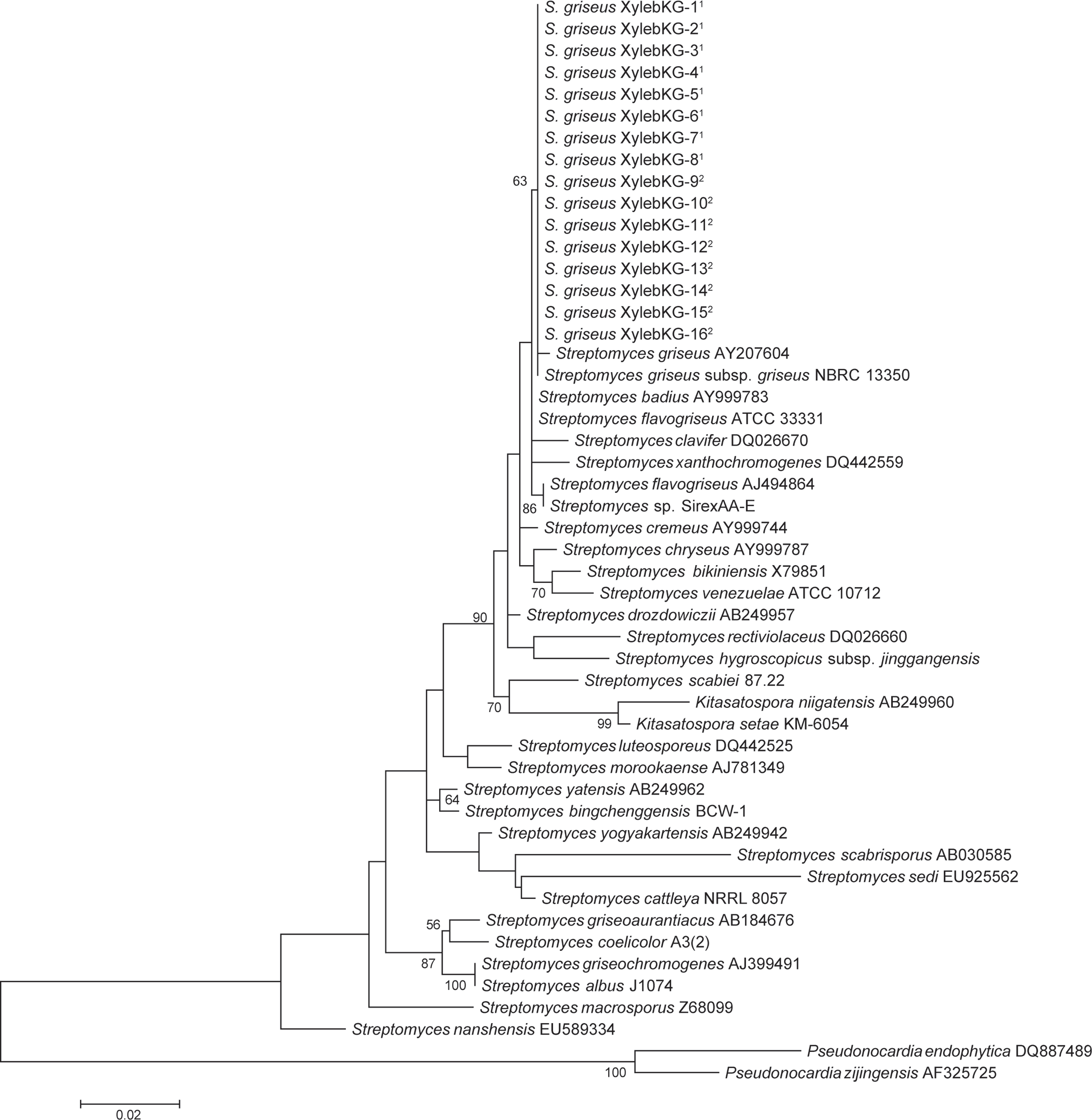
Maximum likelihood 16S phylogeny of the XylebKG-1 clade and its relatives, constructed using MEGA5. Molecular phylogeny based on 1,123 bp of 16S rRNA gene sequence. The Tamura 3-parameter substitution model was used with discrete gamma-distributed rate variation having 5 categories and a proportion of invariable sites, selected by MEGA5 as best fitting the data. The percent node conservation >50% in 500 bootstrap replicates is indicated, and the scale bar indicates the number of substitutions per site. MEGA5's initial heuristic tree search was applied using an initial neighbor-joining tree of pairwise distances estimated using the Maximum Composite Likelihood method. ^1^ - Indicates *X. affinis* origin. ^2^ - Indicates *X. saxesenii* origin.

**Fig. 3.**
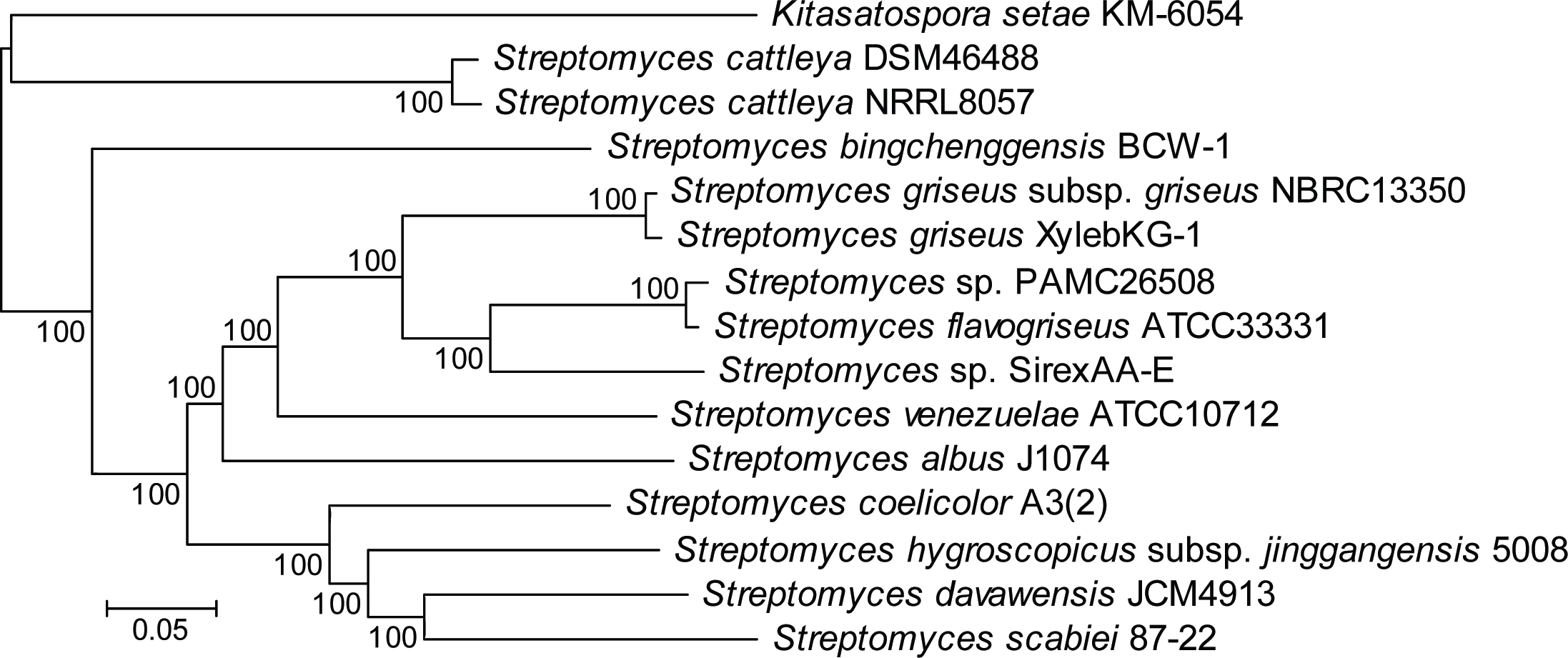
Multilocus phylogeny constructed from 1,364 gene families conserved in all *Streptomyces* genomes analyzed. Alignments were done using MAFFT and the phylogeny generated using RAxML. Numbers above the branches based on 100 rapid bootstraps.

Our work supports a symbiosis between the *S. griseus* XylebKG-1clade and *X. saxesenii* ambrosia beetles. First, strains were consistently isolated having the same culture morphology from nests, larvae, and adults, and a random subset of these had 100 % identical 16S rRNA sequences. Second, Actinobacteria were found to be very abundant within the nest material samples of the investigated *X. saxesenii* strains (approximately 110 *Streptomyces* CFUs per sample). Their recovery rate of 3.4 *Streptomyces* CFUs per adult individual is comparable with other established symbioses, like the *Dendroctonus* bark beetle system (average of 7.7 *Streptomyces* CFUs per individual [27]) or mud daubers (maximum average of 3.1 *Streptomyces* CFUs per individual [33]). Third, *Streptomyces* are vectored by the beetles, likely within their bodies, as artificial medium was sterile and beetles were surface sterilized before being allowed to initiate nests. Fourth, the isolation of the XylebKG-1 Actinobacteria 16S phylotype from *X. affinis* further supports an association with ambrosia beetles, and suggests its potentially wider phylogenetic distribution within these insects.

### Fungal symbionts

Two fungi were consistently isolated from *X. saxesenii* nests. One type was identified as *Raffaelea sulphurea* using a dichotomous key [71] and confirmed by 18S rRNA and EF-α gene sequencing. This fungus has been repeatedly isolated from *X. saxesenii* and is known as the main cultivar of this beetle [4, 9, 71]. The second fungus we isolated was identified as a close relative of the ascomycetous genus *Nectria* based on 18S rRNA and EF-α gene sequences and both BLAST and phylogenetic analyses (Fig. 4). The consistent isolation of this *Nectria* sp. suggests that it is vectored by the ambrosia beetles. *Nectria* species are frequently isolated in low numbers from Scolytine beetles [72], and unpublished 18S rRNA 454-pyrosequencing data from Biedermann *et al*. suggest that they are commonly present in the nests of ambrosia beetles. Given that only *Raffaelea* and *Ambrosiella* species are producing nutritional fruiting structures for feeding ambrosia beetles and *Nectria* spp. are known pathogens of both insects [73] and trees [74], it is likely a parasite of the system.

**Fig. 4.**
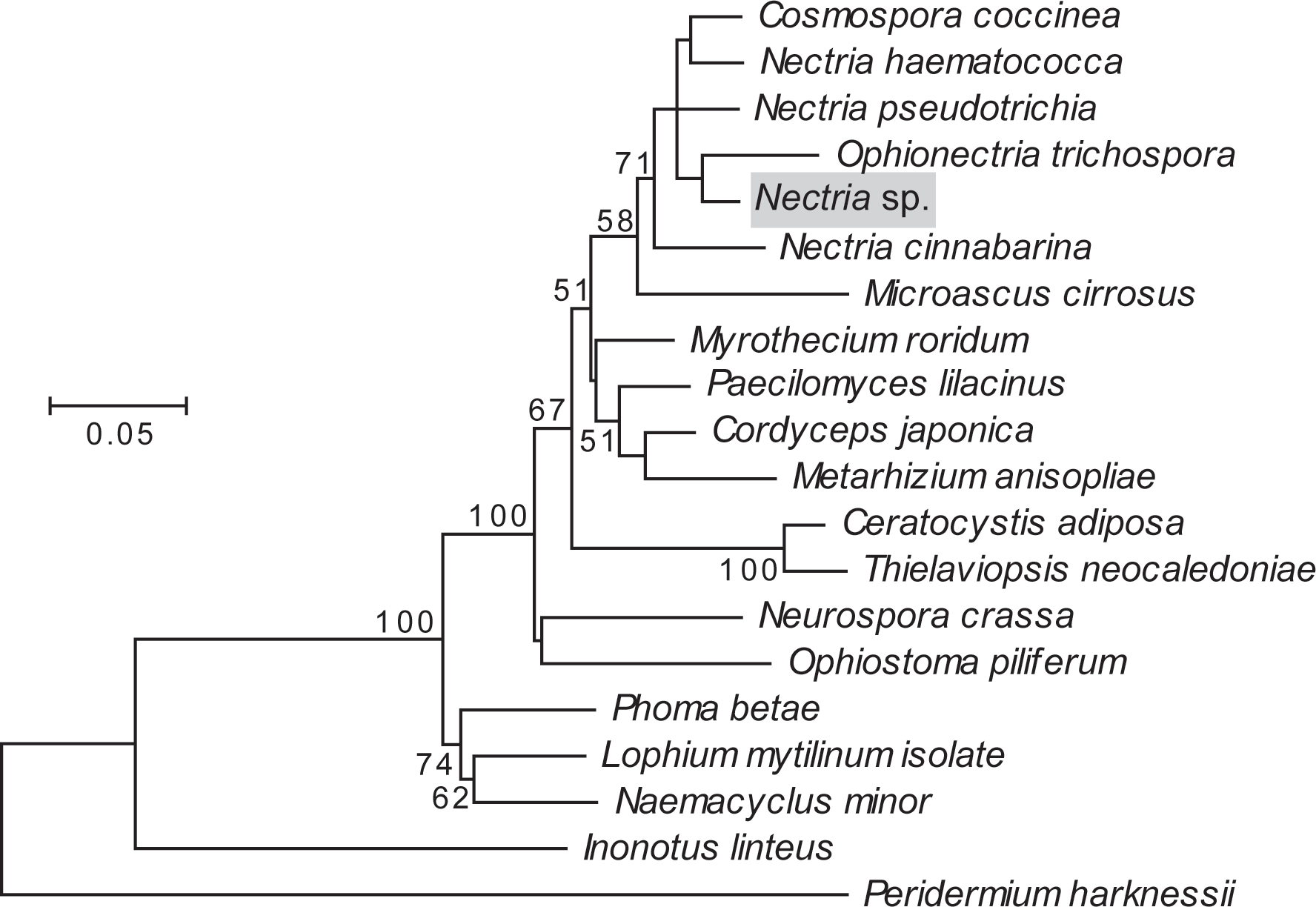
Maximum likelihood phylogeny of the fungal antagonist. highlighted in gray, constructed using concatenated 18S rRNA and EF-α genes in MEGA5. The Tamura-Nei substitution model was used with discrete gamma-distributed rate variation having 5 categories and a proportion of invariable sites, selected by MEGA5 as best fitting the data. The percent node conservation >50% in 500 bootstrap replicates is indicated, and the scale bar indicates the number of substitutions per site. MEGA5's initial heuristic tree search was applied using an initial neighbor-joining tree of pairwise distances estimated using the Maximum Composite Likelihood method.

### The potential of *S. griseus* XylebKG-1 as a defensive mutualist

To explore the potential that *S. griseus* XylebKG-1 function as defensive symbionts of *X. saxesenii*, *S. griseus* XylebKG-1's ability to inhibit the growth of *R. sulphurea* and *Nectria* sp. isolated from this host was examined. Whereas *S. griseus* XylebKG-1 only marginally inhibited the growth of *R. sulphurea* (average zone of inhibition = 0.52 mm; Fig. 5), it significantly inhibited the growth of *Nectria* sp. (average zone of inhibition = 26.2 mm; Fig. 5). The strength of inhibition significantly differed between these fungi (t-test, P = 1.03E-27, n=29 and n=30 for *Nectria* sp. and *R. sulphurea* bioassays, respectively).

**Fig. 5.**
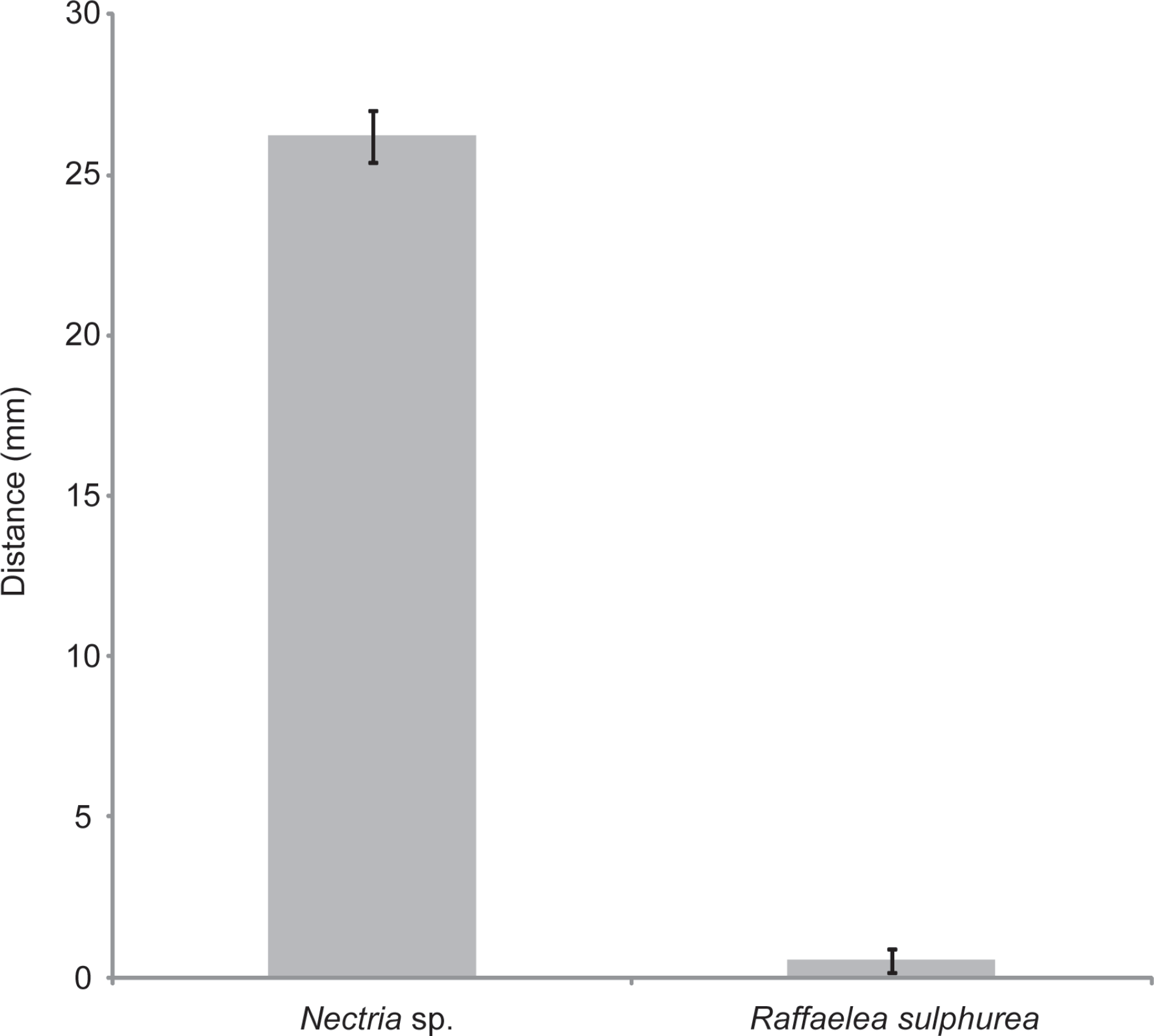
Plate bioassays of *S. griseus* XylebKG-1 versus *Nectria* sp. and *R. sulphurea*. See methods for assay conditions and media used. Average values are shown from 29 and 30 trials (*Nectria* sp. and *R. sulphurea*, respectively), with error bars representing standard error. T-test confirmed significance with P < 0.01.

### Secondary metabolites produced by *S. griseus* XylebKG-1

Cultivation in liquid yeast peptone media led to bioactivity guided isolation of cycloheximide (**1**) using *Nectria* sp. as the indicator organism. HR-ESI-MS provided the molecular formula C_15_H_23_NO_4_; ^1^H and ^13^C NMR data solely matched to the known compound cycloheximide (**1**). In addition, the isolated metabolite showed the same retention time, UV spectrum and ESI-MS pattern as a commercially acquired cycloheximide standard. Furthermore, cycloheximide was produced by *S. griseus* XylebKG-1 cultivated on agar plates of 8 different media (yeast peptone maltose, potato dextrose, oat, soy mannitol, starch glycerol glucose, ISP1, ISP2, and 1187 media), consistent with a robust synthesis of compound cycloheximide under diverse growth conditions. In addition to cycloheximide, two byproducts were isolated by preparative HPLC from a culture of *S. griseus* XylebKG-1 in ISP1 media and identified as naramycin B (**2**) and actiphenol (**3**) [65–67].

**Fig. 6.**
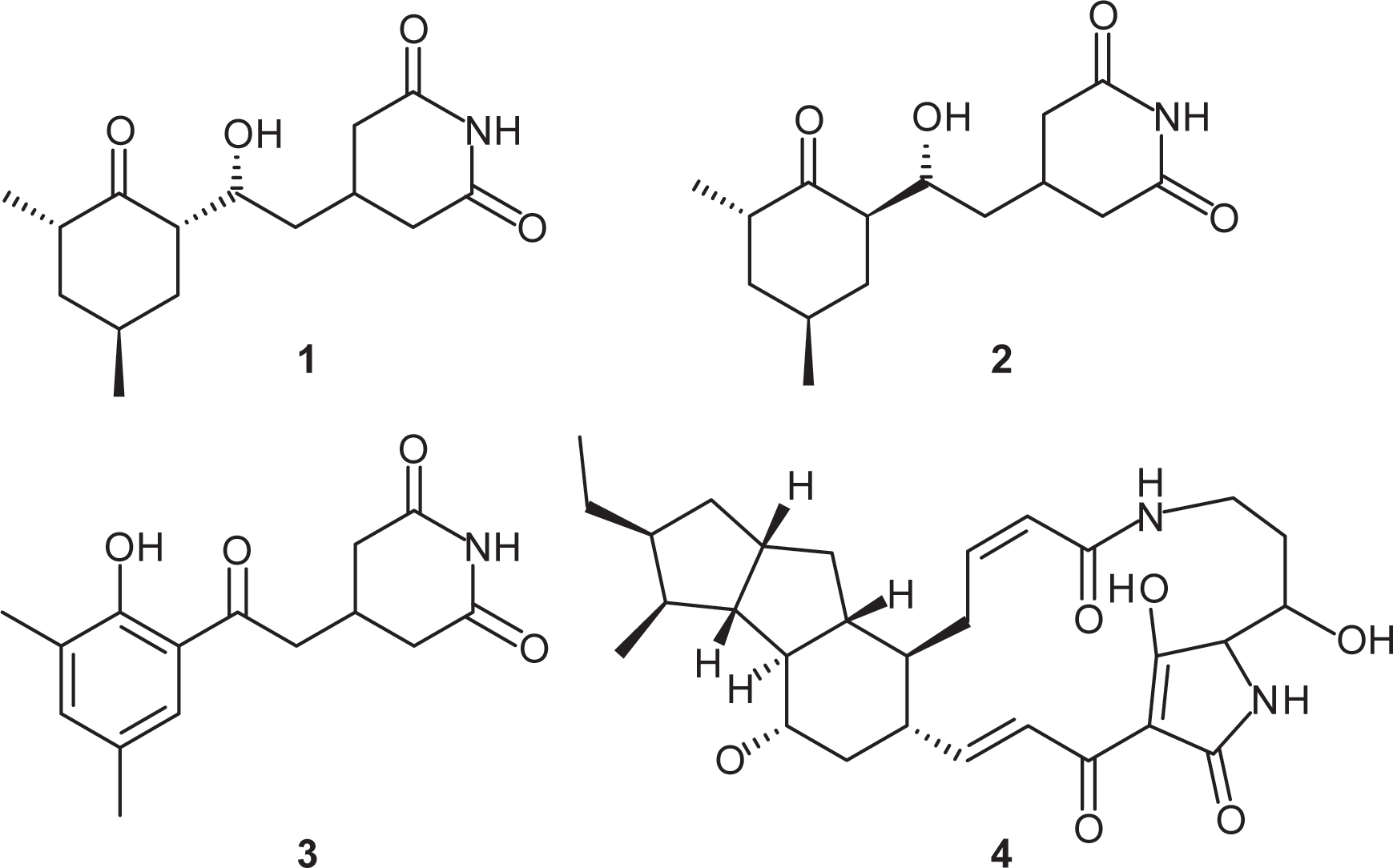
Metabolites isolated from *S. griseus* XylebKG-1. Cycloheximide (**1**); naramycin B (**2**); actiphenol (**3**); dihydromaltophilin (**4**).

Because the alteration of growth media frequently results in a substantially changed metabolite pattern, a switch in growth media can be utilized to explore the metabolic potential of bacterial strains. In the case of *S. griseus* XylebKG-1, cultivation in PD and ISP4 resulted in the biosynthesis of an additional antifungal metabolite. After isolation by preparative HPLC, the molecular formula C_29_H_40_N_2_O_6_ (determined by HR-ESI-MS) and NMR data identified the compound as dihydromaltophilin (**4**) [68].

In plate bioassays, cycloheximide inhibited the isolated *Nectria* sp. (zones of inhibition: 30 μl, 44 mm; 2 μl, 18 mm; 0.2 μg, 9 mm) but not *R. sulphurea* (no inhibition observed). Liquid culture assays confirmed this result, indicating a minimum inhibitory concentration of cycloheximide towards *Nectria* sp. of 0.02 mM. *R. sulphurea* grew in all test conditions (cycloheximide concentration up to 2.7 mM), although it did exhibit slower growth at higher concentrations of cycloheximide (data not shown). Dihydromaltophilin similarly inhibited both *Nectria* sp. and *R. sulphurea* (zones of inhibition: 20 μl, 20 / 22 mm; 2 μl, 12 / 13 mm). Naramycin-B and actiphenol were non-inhibitory under all conditions tested.

A putative cycloheximide biosynthetic gene cluster was identified in the *S. griseus* XylebKG-1 genome using antiSMASH v2.0 [69] based on its homology to the cycloheximide biosynthetic cluster from *Streptomyces* sp. YIM56141 (GenBank accession [JX014302.1]) [70], and its being consistent with retrobiosynthetic logic (data not shown). All proteins predicted in the gene cluster from *Streptomyces* sp. YIM56141 were present in the *S. griseus* XylebKG-1 genome, except that the polyketide synthase *cheE* homolog was annotated as two separate genes in *S. griseus* XylebKG-1 (labeled *cheE1* and *cheE2* in Fig. 7). This entire genomic region is conserved in the genome of *Streptomyces griseus* subsp. *griseus* NBRC13350 [NC_010572.1]. BLAST searches using dihydromaltophilin biosynthetic cluster genes previously identified by Yu *et al*. [75] [EF028635], revealed highly similar protein sequences with the top blast hits of the first six genes (expect values: 0.0, 0.0, 0.0, 0.0, 0.0, and 2e-105) in the cluster being in the same relative positions. Similar to the pine beetle symbiont *Streptomyces* sp. SPB78 in which no homologs were found for ferredoxin-like and arginase proteins [76], BLAST indicated top hits for these genes in separate areas of the genome with much higher expect values (expect values: 8e-009 & 2e-010 respectively). The same pattern of results appeared in a BLAST search of the same cluster fragment genes against *S. griseus* subsp. *griseus* NRBC13350 [NC_010572.1]. Namely, the top BLAST hits for the first six genes had a conserved genomic order and extremely small expect values (0.0, 0.0, 0.0, 0.0, 0.0, and 2e-105) while the last two genes (ferredoxin-like and arginase) had much higher expect values (2e-010 & 8e-009) and were found in other parts of the genome. Evolutionary conservation of these clusters from before the adaptation of *S. griseus* XylebKG-1 as a symbiont of ambrosia beetles suggests the potential for similar and/or complementary regulation and activity of the metabolites that they produce.

**Fig. 7.**
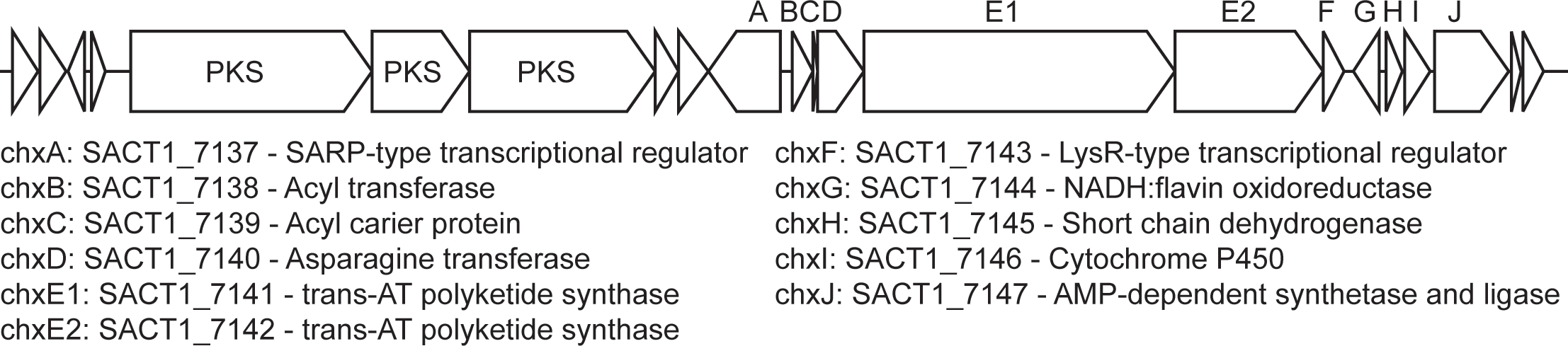
Putative cycloheximide biosynthetic gene cluster in *S. griseus* XylebKG-1, predicted using antiSMASH v2.0, homology to a cycloheximide biosynthetic gene cluster from *Streptomyces* sp. YIM56141, and retrobiosynthetic logic. Letters above the cluster indicate gene names, with corresponding locus names and annotations indicated below. An upstream biosynthetic gene cluster predicted to be unrelated to cycloheximide biosynthesis is also shown, with its polyketide synthase (PKS) genes indicated.

Our findings are consistent with *S. griseus* XylebKG-1 being a potential defensive symbiont of ambrosia beetles. Cycloheximide's specific inhibition of the antagonist *Nectria* sp., but not the mutualist *R. sulphurea*, supports a defensive role for its production by XylebKG. This parallels similar results obtained for fungus-growing ants and *Dendroctonus frontalis*, where their associated Actinobacteria inhibited the growth of a fungal parasite and not the fungal mutualist [26, 27]. Cycloheximide inhibits protein synthesis in eukaryotic cells and as such is toxic to most eurkaryotes [77], including fungi. Interestingly, species in the fungal order Ophiostomatotales (including *Raffaelea* spp.), are known to largely be resistant to cycloheximide [28, 78]. Although not determined here, *X. affinis* also cultivates a cycloheximide-insensitive *Raffaelea* sp. as food [47]. Two different ambrosia beetle species associating with two different cycloheximide-insensitive *Raffaelea* spp. mutualists as well as the cycloheximide-producing *S. griseus* XylebKG-1 further supports the role of these bacteria as defensive symbionts, as evolutionarily stable relationships are expected to promote such complementarity. In contrast, dihydromaltophilin production inhibits the growth of both *Nectria* sp. and *R. sulphurea*. The production of dihydromaltophilin under only a few growth conditions could suggest that it does not have an active role in the ambrosia beetle system, but rather is a remnant from before *S. griseus* XylebKG-1 became associated with these beetles. In this regard it is worth noting that dihydromaltophilin analogs were found at low expression levels in the *Dendroctonus frontalis* system [76]. Alternatively, dihydromaltophilin production may be regulated to avoid inhibition of *R. sulphurea*, or selected for activity versus other organisms not considered in this study.

## Conclusion

In this study we consistently isolated a single *Streptomyces* morphotype and phylotype from both *X. saxesenii* and *X. affinis* that inhibited the growth of the parasitic fungus *Nectria* sp., but not the mutualistic *R. sulphurea*, likely via the production of cycloheximide. Its ubiquity suggests that XylebKG-1 may be a defensive mutualist of these ambrosia beetles that inhibits the growth of all but a few fungi, including its mutualistic fungal food source. Future studies should include natural nests collected from a wider range of species and geographies to establish the breadth and prevalence of XylebKG-1 in bark and ambrosia beetle nests [28]. These studies should also include greater phylogenetic power as 16S r RNA gene analyses is not sufficient to resolve species within the *S. griseus* clade [79]. Cycloheximide should be assayed *in vivo* to confirm the relevance of its *in vitro* activity, and any other compounds also produced determined *in vivo* (e.g., those produced by the biosynthetic gene cluster adjacent to the cycloheximide cluster), if they exist. Furthermore, the presence and activity of XylebKG-1 may also vary during beetle development, e.g., cycloheximide may be used to clear new nests of contaminating fungus in preparation for the agricultural symbiont. The antibiosis of cycloheximide includes a large non-specific range of fungi and as such other fungal symbionts, aside from *Nectria* sp., may also be inhibited. We have identified a putative defensive Actinobaterium and an antagonistic fungal symbiont in two ambrosia beetles, potentially expanding the interactions from bipartite to quadripartite.

**S1 Methods. Additional details concerning analytical chemistry methods.**

**S2 Fig1. Updated Maximum likelihood 16S phylogeny of the XylebKG-1 clade and its relatives**

**S3 Fig2. ^1^H spectrum of cycloheximide (CD_3_OD).**

**S4 Fig3. ^1^H spectrum of naramycin B (CD_3_OD)**

**S5 Fig4. ^1^H spectrum of dihydromaltophilin (CD_3_OD)**

