## Supplementary Methods and Figures for "Cycloheximide-Producing *Streptomyces* Associated with *Xyleborinus saxesenii* and *Xyleborus affinis* Fungus-Farming Ambrosia Beetles"

**General Experimantal Procedures.** $^{1}H$ and two dimensional NMR spectra were acquired using a Varian Inova 600 MHz spectrometer, $^{13}C$ spectra were acquired using a Varian Inova 150 MHz spectrometer. HPLC/MS analysis was performed on an Agilent 1200 Series HPLC / 6130 Series mass spectrometer. High resolution spectra were obtained on a Waters Micromass Q-Tof Ultima ESI-TOF mass spectrometer.

**Cultivation of *Streptomyces* sp. Xs1.** For strain maintenance, *Streptomyces* sp. Xs1 was cultivated on potato dextrose (PD) agar plates containing per liter 4 g potato starch, 20 g dextrose for 5 - 10 days. As a preculture, 3x 85 mL yeast peptone maltose (YPM) media containing per liter 2 g yeast extract, 2g bactopeptone, 4g D-mannitol in 500 mL Erlenmeyer flasks were inoculated with an about 1 $cm^2$ piece of an agar plate overgrown well with *Streptomyces* sp. Xs1 and incubated at 28 °C and 250 rpm on a rotating shaker for 48 h. 8x 1 L yeast peptone media in 4 L Erlenmeyer flasks without baffles were inoculated with 25 mL of the described preculture each and incubated for 72 h at 28 °C and 250 rpm. For harvesting, the culture was centrifuged (7000 rpm, 30 min) and the supernatant and mycelium were worked up separately.

**Isolation of cycloheximide.** The supernatant was adjusted to pH = 6 and extracted twice with an equal amount of ethyl acetate. After evaporation in vacuo the residue was resuspended in 2 mL $MeOH/H_2O$ (8:2). The mycelium was lyophilized and extracted with 50 mL aceton and 50 mL methanol. After evaporation in vacuo the crude extract was resuspended in 2 mL methanol. Crude extracts of the supernatant and the mycelium were tested for inhibition of *Melanospora fallax* at various concentrations; only the extract of the supernatant showed significant bioactivity. The crude extract was loaded on a 2 g pre-packet $C_{18}$ Sepak resin and fractionated by eluting with a gradient from pure water to pure methanol. The flow through and the 10 % methanol fraction were most active in inhibiting the growth of the opportunistic fungus *Melanospora fallax*. These fractions were combined and fractionated by gel chromatography using Sephadex LH-20 with methanol as mobile phase (column 60 x 2.5 cm). Active fractions were combined and subsequently purified by reversed-phase HPLC (Agilent 1100 Series HPLC system, Supelco Discovery HS C18 column, 250 x 10 mm, 2 mL/min). HPLC conditions used: 2 min 80 % A, 20 % B in 28 min to 100 % B (A: water, B: methanol). Most active against *Melanospora fallax* was the fraction which eluted from 17.5 and 18 min. It was further analyzed by HPLC/MS and contained 2.7 mg of cycloheximide:

ESI-MS: $[M+Na]^+$ 304.1, $[M+H]^+$ 282.1, $[M-H]^-$ 280.1; meassured HR-ESI-MS 282.1718, calculated HR-ESI-MS $[M+H]^+$ 282.1700 ($C_{15}H_{24}NO_4$); UV: 215 nm; OD $[\alpha]_D^{20}$ -0.514 (c = 1.7 g/L in MeOH).

**Comparision to commercial cycloheximide standard.** From *Streptomyces* sp. Xs1 isolated cycloheximide was compared to a commercial standard with $^{1}H$ and $^{13}C$ NMR spectroscopy and HPLC-MS analysis regarding retention time, UV spectrum and electron spray mass spectrum.

**OSMAC Screening.** *Streptomyces* sp. Xs1 was cultivated on agar plates (300 mL) of YPM, PD, oat media (per liter 20 g oat meal, 2.5 mL trace element solution; trace element sol.: 3 g $CaCl_2 \cdot 2\,H_2O$, 1 g Fe(III)-citrate, 0.2 g $MnSO_4$, 0.1 g $ZnCl_2$, 25 mg $CuSO_4 \cdot 5\,H_2O$, 20 mg $Na_2B_4O_7 \cdot 10\,H_2O$, 4 mg $CoCl_2$, 10 mg $Na_2MoO_4 \cdot 2\,H_2O$), soy mannitol media (per liter 20 g soy meal, 20 g mannitol), starch-glucose-glycerol media (per liter 10 g glucose, 10 g glycerol, 10 g starch, 2.5 mL cornsteep liquor, 5 g casein-peptone, 2 g yeast extrakt, 1 g NaCl, 3 g $CaCO_3$), ISP1 media (per liter 5 g pancreatic digest of casein, yeast extract 3 g), ISP2 media (per liter 4 g yeast extract, 10 g malt extract, 4 g dextrose), 1187 meida (per liter 10 g starch, 2 g $(NH_4)_2SO_4$, 1 g $K_2HPO_4$, 1 g $MgSO_4 \cdot 7\,H_2O$, 1 g NaCl, 2 g $CaCO_3$, 5 mL trace element solution described above) for 7 days at 30 °C. For harvesting the plates were extracted with ethyl acetate. Dihydomaltophilin was isolated from extracts of PD and 1187 cultivations by preparative HPLC (gradient 50 % to 100 % acetonitrile in 25 min, column Phenomenex Luna C18 250x21mm), (gradient 80 % to 100 % acetonitrile in 25 min, column Phenomenex Luna C18 250x21mm) and semi-preparative HPLC (gradient 35 % to 50 % acetonitrile in 25 min, column Supelco C18 250x8mm).

ESI-MS: $[M+H]^+$ 513.3, $[M-H]^-$ 511.3; meassured HR-ESI-MS 513.2964, calculated HR-ESI-MS $[M+H]^+$ 513.2965 ($C_{29}H_{41}N_2O_6$); UV: 215, 320 nm.

Naramycin B (0.7 mg) was isolated from the ISP1 cultivation by semipreparative HPLC (gradient 22 % to 40 % acetonitrile in 25 min, Supelco Discovery HS C18 column, 250 x 10 m).

ESI-MS: $[M+Na]^+$ 304.1, $[M+H]^+$ 282.1, $[M-H]^-$ 280.1; UV: 215 nm; OD $[\alpha]_D^{20}$ +0.015 (c = 0.7 g/L in MeOH).

Actiphenol (1.6 mg) was isolated from the 1187 cultivation by preparative HPLC (gradient 65 % to 100 % methanol in 25 min, column Phenomenex Luna C18 250x21mm).

ESI-MS: $[M+Na]^+$ 298.0, $[M+H]^+$ 276.2, $[M-H]^-$ 274.1; UV: 270, 350 nm.

**Cultivation of *Streptomyces* sp. Xa1.** Streptomyces sp. Xa1 was cultivated in 8 L YPM medium for 3 days in under the same conditions as sp. Xs1. The active metabolite (74.4 mg) could be isolated by two rounds of gel chromatography (sephadex LH-20, 1. methanol, 2. acetone) and two steps of reverse phase HPLC (1. gradient 20% to 100% methanol in 30 min, column Supelco Discovery C18; 2. gradient 30% to 60% acetonitril in 30 min, column Alltech Alltima C18).

**Liquid antifungal assay.** *Melanospora fallax* was grown in yeast peptone media for three days. The culture was diluted 1:1000 with fresh yeast peptone media and 200µL were transferred into each well of a 96-well plate and various concentration of the test substances were applied. The plates were incubated for 72 h at 30 °C and 20 mL almarBlue (Soretec Ltd.) was added. The fluorescence was measured with excitation at 540 nm and emission at 590 nm by a SpectraMax M5® Plate Reader after 12 h inoculation at 30 °C.

**Plate antifungal assay.** *Melanospora fallax* and *Ambrosiella sulfurea* were grown in 20 mL potato dextrose media for 7 respectively 21 days at 30 °C and 250 rpm. 1 ml of these cultures were put on the top of potato dextrose agar plates and allowed to dry under sterile conditions. Paper disks (6 mm diameter) were soaked with solutions of 30 µg, 2 µg respectively 0.2 µg cycloheximide in methanol. The disks were dried and applied to the surface of the agar plates.

S2. Figure 1

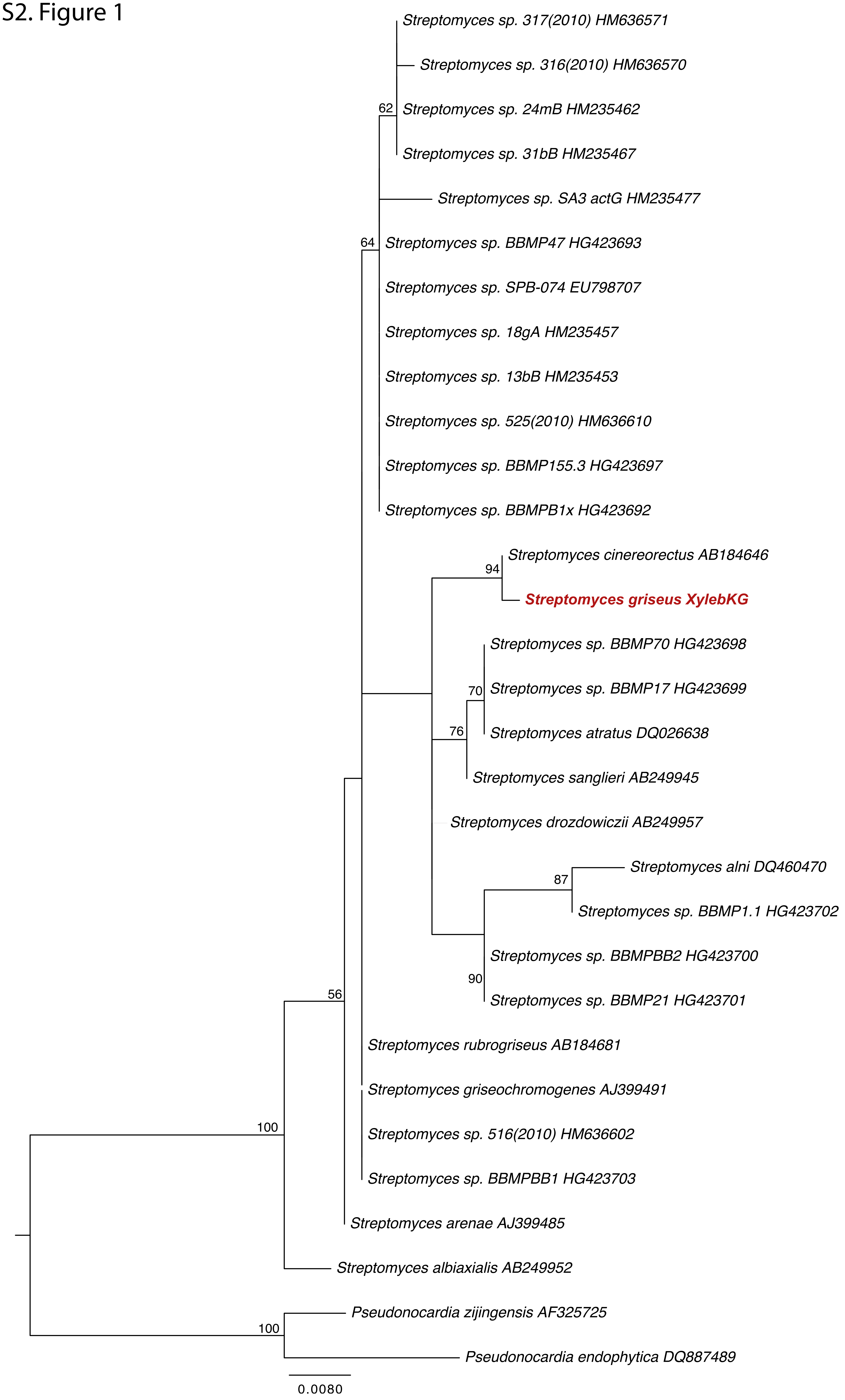
Streptomyces sp. 317(2010) HM636571
Streptomyces sp. 316(2010) HM636570
62
Streptomyces sp. 24mB HM235462
Streptomyces sp. 31bB HM235467
Streptomyces sp. SA3 actG HM235477
64
Streptomyces sp. BBMP47 HG423693
Streptomyces sp. SPB-074 EU798707
Streptomyces sp. 18gA HM235457
Streptomyces sp. 13bB HM235453
Streptomyces sp. 525(2010) HM636610
Streptomyces sp. BBMP155.3 HG423697
Streptomyces sp. BBMPB1x HG423692
Streptomyces cinereorectus AB184646
94
Streptomyces griseus XylebKG
Streptomyces sp. BBMP70 HG423698
70
Streptomyces sp. BBMP17 HG423699
76
Streptomyces atratus DQ026638
Streptomyces sanglieri AB249945
Streptomyces drozdowiczii AB249957
Streptomyces alni DQ460470
87
Streptomyces sp. BBMP1.1 HG423702
Streptomyces sp. BBMPBB2 HG423700
90
Streptomyces sp. BBMP21 HG423701
56
Streptomyces rubrogriseus AB184681
Streptomyces griseochromogenes AJ399491
100
Streptomyces sp. 516(2010) HM636602
Streptomyces sp. BBMPBB1 HG423703
Streptomyces arenae AJ399485
Streptomyces albiaxialis AB249952
Pseudonocardia zijingensis AF325725
100
Pseudonocardia endophytica DQ887489
0.0080

S3. Figure 2

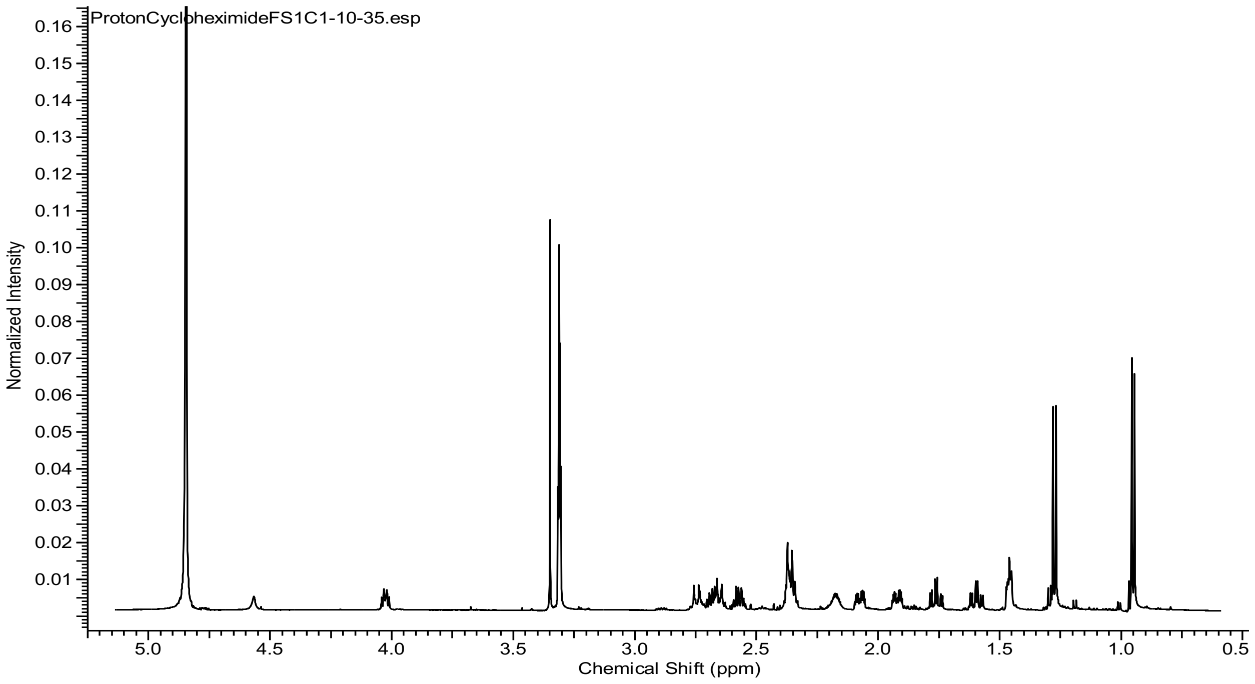
ProtonCycloheximideFS1C1-10-35.esp
Normalized Intensity
0.16
0.15
0.14
0.13
0.12
0.11
0.10
0.09
0.08
0.07
0.06
0.05
0.04
0.03
0.02
0.01
5.0
4.5
4.0
3.5
3.0
2.5
2.0
1.5
1.0
0.5
Chemical Shift (ppm)

S4. Figure 3

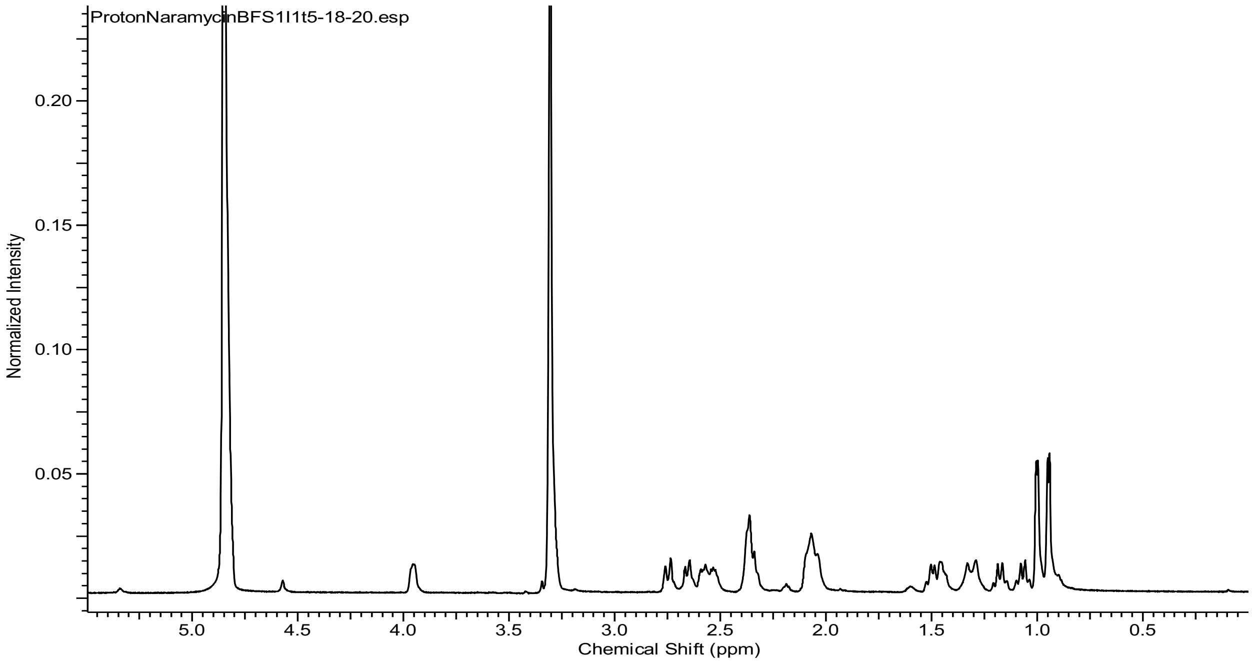
ProtonNaramycinBFS1I1t5-18-20.esp
Normalized Intensity
0.20
0.15
0.10
0.05
5.0
4.5
4.0
3.5
3.0
2.5
2.0
1.5
1.0
0.5
Chemical Shift (ppm)

S4. Figure 4

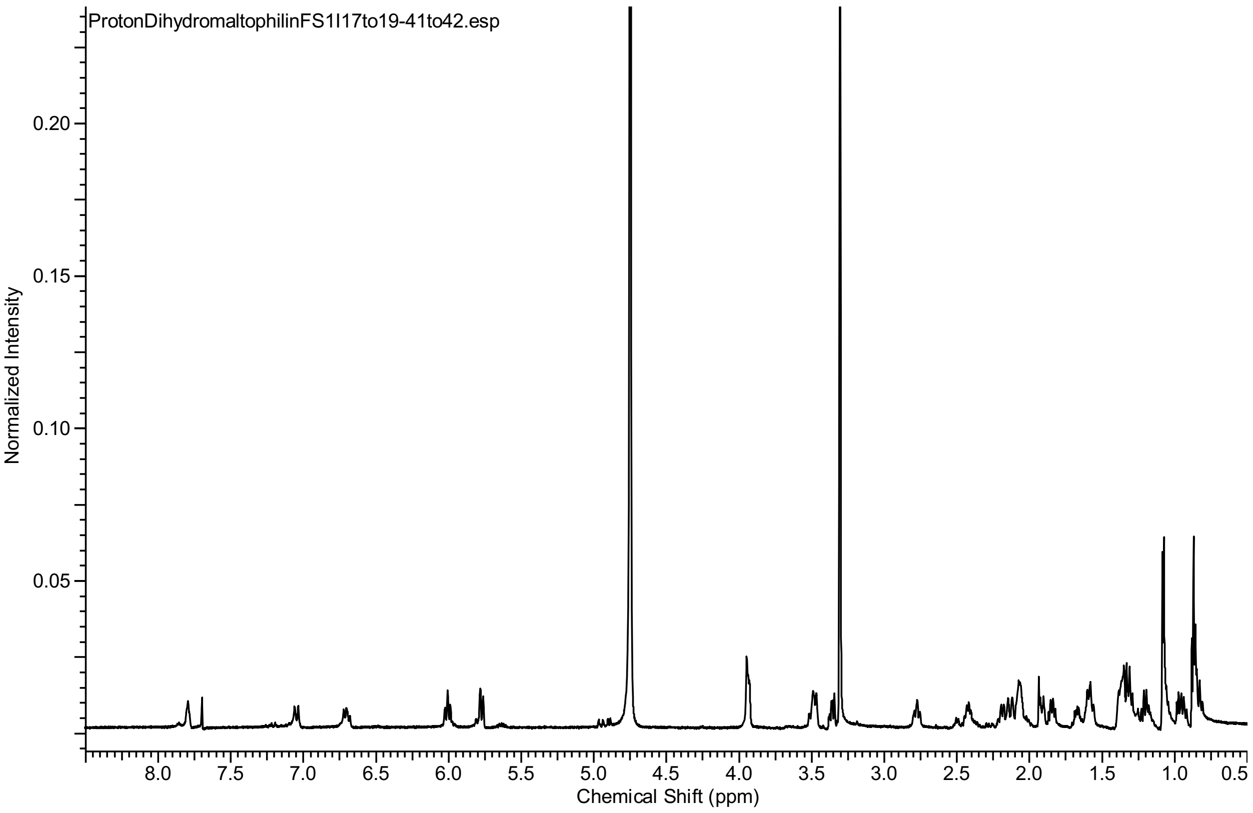
ProtonDihydromaltophilinFS1I17to19-41to42.esp
Normalized Intensity
0.20
0.15
0.10
0.05
8.0
7.5
7.0
6.5
6.0
5.5
5.0
4.5
4.0
3.5
3.0
2.5
2.0
1.5
1.0
0.5
Chemical Shift (ppm)
